# Structural Topology-based Electrostatic Model (STEM) Reveals Ion-Coordination Exchange as a Driver of RNA Folding Dynamics

**DOI:** 10.64898/2026.06.27.734987

**Authors:** Avijit Mainan, Akhilesh Jaiswar, José N Onuchic, Karissa Y. Sanbonmatsu, Susmita Roy

## Abstract

RNA is a highly charged polyelectrolyte whose folding into functional architectures depends on an ionic atmosphere that screens strong electrostatic repulsion along the phosphate backbone. Whereas monovalent ions primarily stabilize secondary structure, divalent magnesium (Mg^2+^) drives tertiary folding often via site-specific and adopting various dynamic coordination modes. Current RNA structure-prediction frameworks rely largely on static direct-contact information, overlooking ion-mediated interactions and the dynamic exchange between distinct coordination modes—particularly the dynamic exchange between direct (inner) and solvent-separated (outer-sphere) Mg^2+^-phosphate coordination that often controls RNA’s conformational transition. Here, we introduce the Structural-based Electrostatic Model (STEM), a hybrid implicit–explicit framework that explicitly captures how the dynamic exchange between distinct ion-coordination modes dictates folding pathways. STEM combines explicit Mg^2+^ ions to resolve site-specific interactions with implicit K^+^ ions to describe counter-ion condensation mediated electrostatic screening through generalized Manning counter-ion condensation model, enabling computationally efficient exploration of RNA folding landscapes. The model accurately reproduces crystallographic ion-binding sites, experimental preferential ion-interaction coefficients, and Small-Angle X-ray Scattering (SAXS)-derived radii of gyration across diverse RNA systems. Applied to a 58-nt rRNA fragment, STEM reveals that folding from an intermediate to the native state is driven by a chelated Mg^2+^-mediated tertiary contact and captures the resulting coordination-dependent conformational breathing. By shifting the paradigm from static direct-contact descriptions to ion-mediated dynamic interactions, STEM provides a physically grounded framework for predicting dynamic ensembles of RNA structures, resolving their folding free-energy landscapes, and elucidating the mechanisms of RNA folding and function beyond native conformations across physiological salt conditions.

**GRAPHICAL ABSTRACT:** 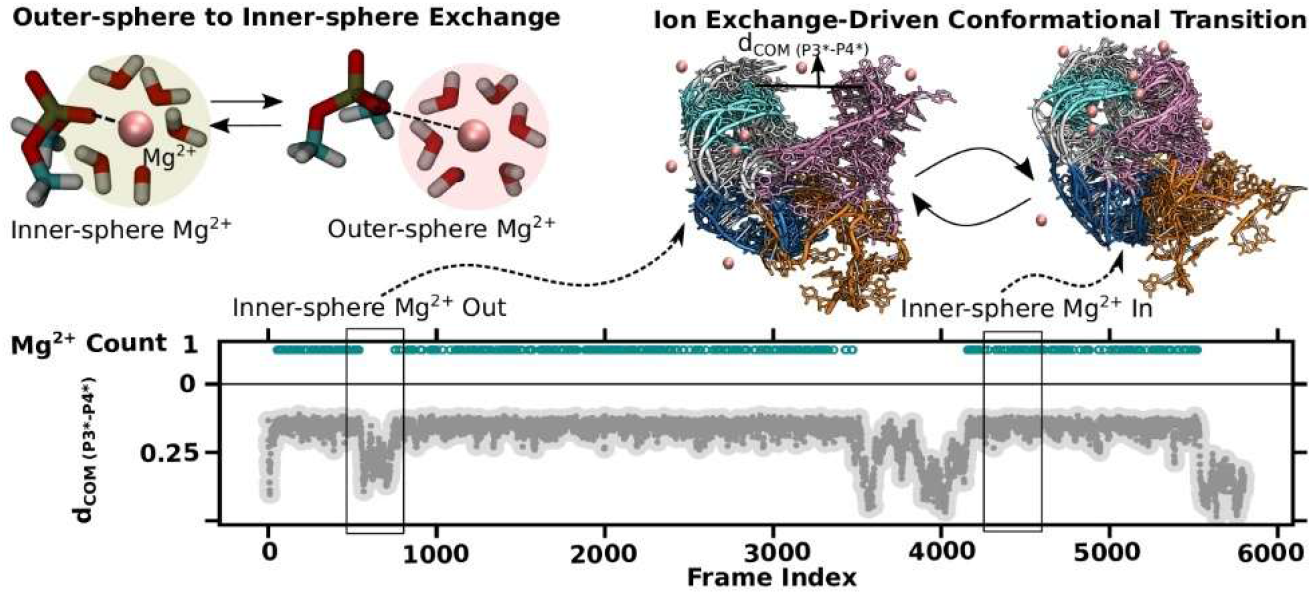

## INTRODUCTION

As water shapes stone through persistent yet unseen forces, ions sculpt the folding landscape of RNA. Despite carrying strong electrostatic repulsion along its phosphodiester backbone, RNA folds into compact and functional architectures through the stabilizing influence of a surrounding ionic atmosphere. Under physiological conditions, this atmosphere comprises monovalent (e.g., Na^+^, K^+^) and divalent (e.g., Mg^2+^, Ca^2+^) cations together with water molecules. Beyond simple charge neutralization, these ions form condensed ionic layers that provide long-range electrostatic stabilization essential for maintaining compact and biologically active RNA conformations. Experimental and computational studies have shown that divalent Mg^2+^ ions are the principal determinants of RNA tertiary folding through preferential interactions with phosphate groups (1–8) whereas monovalent cations, particularly Na^+^ and K^+^ (9, 10), predominantly stabilize secondary structural elements by electrostatically screening the negatively charged backbone. Remarkably, although intracellular K^+^ concentrations (∼100–300 mM) exceed physiological Mg^2+^ concentrations (∼0.5–1.5 mM) by two orders of magnitude(11, 12), Mg^2+^ exerts a disproportionately strong influence on RNA compaction and tertiary organization owing to its high charge density and unique ability to engage in site-specific coordination. This observation raises a fundamental question: how can a relatively small population of Mg^2+^ ions exert such profound control over RNA structure and dynamics?

The answer appears to lie not only in the presence of Mg^2+^ ions but also in the manner by which they interact with RNA. The ionic atmosphere surrounding RNA is far from a passive cloud of charge-compensating ions; rather, it is a dynamic environment whose constituents continuously reorganize as RNA explores its conformational landscape. To conceptualize this complexity, Draper and co-workers classified the RNA ion atmosphere into two broad categories: diffuse ions, which remain largely hydrated in solution, and chelated ions, which establish direct interactions with negatively charged RNA groups(1). Subsequent studies refined this picture by identifying outer-sphere Mg^2+^ ions that remain separated from RNA by a single hydration layer yet occupy preferred locations around the molecule, thereby bridging the gap between fully diffuse and directly coordinated ions(10, 13, 14). Together, these observations reveal that Mg^2+^ ions populate a continuum of coordination states rather than a simple bound–unbound equilibrium.

This distinction is particularly important because different coordination modes appear to exert fundamentally different effects on RNA structure. Chelated Mg^2+^ ions can simultaneously interact with distant phosphate groups, stabilizing tertiary contacts that would otherwise be destabilized by electrostatic repulsion(15). For example, crystal structures of Group I introns revealed strategically positioned Mg^2+^ ions that stabilize the catalytic core and are indispensable for ribozyme activity (16, 17). Yet such structural snapshots provide only static views of a fundamentally dynamic process and do not reveal whether transitions between coordination states actively participate in RNA folding.

Intriguing clues emerge from folding experiments. Draper and colleagues demonstrated that an RNA intermediate (I) state is stabilized primarily by monovalent ions and outer-sphere Mg^2+^ interactions, whereas formation of the native (N) state coincides with the appearance of a chelated inner-sphere Mg^2+^ ion(18). Thermodynamic analyses further showed that this chelated ion substantially stabilizes the native fold relative to the intermediate ensemble. These findings suggest a compelling mechanistic picture: rather than serving merely as static structural cofactors, Mg^2+^ ions may actively drive folding through dynamic transitions between outer-sphere and inner-sphere coordination states. In this view, RNA folding emerges from a continual interplay between conformational rearrangements of the molecule and reorganization of its ionic environment, with ion-coordination exchange reshaping the underlying free-energy landscape.

Testing this hypothesis remains challenging because RNA folding and Mg^2+^-coordination exchange span multiple spatial and temporal scales. Experimental techniques provide valuable structural and thermodynamic insights but offer limited access to the microscopic pathways connecting distinct coordination states. Likewise, conventional all-atom molecular dynamics simulations remain computationally prohibitive for capturing the long-timescale fluctuations and rare ion-exchange events associated with RNA folding. Consequently, a variety of theoretical and computational frameworks have been developed to describe RNA electrostatics. Continuum approaches based on the Poisson–Boltzmann equation successfully capture mean electrostatic screening, whereas more advanced methods such as size-modified Poisson– Boltzmann, 3D-RISM, and tightly bound ion (TBI) models incorporate ion size, solvent structure, and ion–ion correlations(5, 19–30). Although these approaches have significantly advanced our understanding of RNA ion atmospheres(31, 32), they generally treat ion coordination as a static property and do not explicitly account for dynamic exchange between outer-sphere and inner-sphere Mg^2+^ states. Previous models incorporated the energetic effects of inner- and outer-sphere Mg^2+^ interactions, but generally treated them as static interaction channels or equilibrium potentials(31). STEM explicitly links the dynamic exchange between coordination states to RNA conformational dynamics and folding transitions, allowing ion exchange itself to emerge as a mechanistic variable along the folding landscape. (30)

Motivated by this gap, we sought to develop a framework that bridges ion-coordination exchange and RNA conformational dynamics within a single computationally tractable model. Here, we introduce the Structural Topology-based Electrostatic Model (STEM). To achieve both physical realism and computational efficiency, STEM combines implicit monovalent-ion screening through Generalized Manning Counterion Condensation theory(33) inspired by Classical Manning Condensation Theory(34) with explicit Mg^2+^ interactions and, crucially, allows dynamic exchange between outer-sphere and inner-sphere coordination states. By treating ion coordination as an active participant rather than a passive consequence of RNA folding, STEM provides a physically grounded framework for investigating how a small number of strategically positioned Mg^2+^ ions reshape RNA conformational ensembles, folding free-energy landscapes, and folding pathways.

## Model

### Development of the Structural Topology-based Electrostatic Model (STEM)

To quantitatively investigate how Mg^2+^-coordination exchange influences RNA conformational dynamics, we developed the Structural Topology-based Electrostatic Model (STEM), a hybrid electrostatic framework that integrates RNA structural topology, counterion condensation, and dynamic ion-coordination exchange within a computationally tractable formalism. STEM builds upon Generalized Manning Counterion Condensation (GMCC) theory(33) with explicit divalent-ion interactions. Under physiological conditions, RNA exists in a highly concentrated monovalent ionic environment (>100 mM), where a substantial fraction of counterions accumulates within the RNA ion atmosphere and contributes primarily to electrostatic screening and fold stabilization. To account for this effect, STEM employs the GMCC framework to represent monovalent ions implicitly while treating Mg^2+^ ions explicitly because of their central role in site-specific coordination and tertiary-structure stabilization. This hybrid implicit–explicit treatment preserves the essential physics of the RNA ion atmosphere while accurately describing Mg^2+^-mediated interactions and their dynamic exchange between coordination states. By avoiding the explicit representation of the large population of highly dynamic monovalent ions and solvent molecules, STEM achieves substantial computational efficiency, enabling the investigation of large-scale conformational fluctuations, folding transitions, and long-timescale ion-coupled dynamics that are otherwise inaccessible to conventional all-atom simulations.

STEM Hamiltonian consists of three physically distinct contributions,

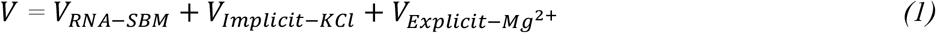

Where *V*_*RNA–SBM*_ is the structure-based model (SBM) potential for RNA(35), *V*_*Implicit−KCl*_ accounts for the collective electrostatic screening arising from the monovalent ionic atmosphere, and 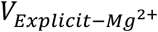 captures Mg^2+^-mediated interactions together with dynamic exchange between outer-sphere and inner-sphere coordination states.

The RNA folding landscape is represented using an all-heavy-atom structure-based model (SBM) previously developed for RNA systems(35–37). The potential consists of local bonded interactions (bonds, angles, and dihedrals) together with native-contact attractive interactions and non-native repulsive interactions. Such formulation has been widely used and validated for describing conformational fluctuations and folding dynamics in RNA and RNA-protein assemblies(35, 38). A complete description of the SBM potential is provided in our early work(8).

To account for the strong accumulation of monovalent counterions around RNA under physiological salt conditions, STEM employs the Generalized Manning Counterion Condensation (GMCC) framework(33). Within this approach, condensed K^+^ ions are represented implicitly as Gaussian charge distributions surrounding phosphate groups, thereby capturing heterogeneous electrostatic screening without explicitly simulating the large population of rapidly exchanging monovalent ions. The detailed derivation of the GMCC formalism and the associated mixing free-energy terms are provided in our early work (31).

### Explicit Mg^2+^ Treatment and Ion-Coordination Exchange

The principal advance of STEM lies in the explicit treatment of Mg^2+^ ions and the incorporation of dynamic exchange between outer-sphere and inner-sphere coordination states. Because Mg^2+^-mediated tertiary interactions are highly localized and cannot be adequately described by continuum electrostatics, we first quantified the underlying Mg^2+^-phosphate free-energy landscape using explicit-solvent molecular dynamics simulations.

### Quantifying Mg^2+^ –Phosphate Free-Energy Landscape

To incorporate dynamic exchange between outer-sphere and inner-sphere Mg^2+^ coordination into the STEM Hamiltonian, two complementary explicit-solvent approaches were employed: (i) The first involved direct PMF calculations using umbrella sampling(39) of a simple model system coordinated with one Mg^2+^ ion; (ii) the second derived effective free-energy profiles through Boltzmann inversion of Mg^2+^-phosphate radial distribution functions (RDFs) obtained from a number of equilibrated RNA simulations.

For the umbrella-sampling calculations, a simple model, Mg^2+^-dimethyl phosphate (DMP) system was chosen to isolate the fundamental Mg^2+^-phosphate interaction from the system-specific topological complexity of folded RNA. The free-energy profile was computed using the AMBER force field with Mg^2+^ parameters optimized for RNA coordination(40–42). Details of the simulation protocol are provided in the Supporting Information. The resulting PMF exhibits two well-defined minima separated by a substantial free-energy barrier, corresponding to inner-sphere and outer-sphere Mg^2+^ coordination states (**Figure 1A**). Notably, the free-energy difference between these states is relatively small, whereas the intervening barrier is sufficiently large to generate distinct metastable coordination modes. Consistent with this picture, we recently benchmarked the Mg^2+^-phosphate exchange kinetics against ^25^Mg NMR measurements(43).

**Figure 1:**
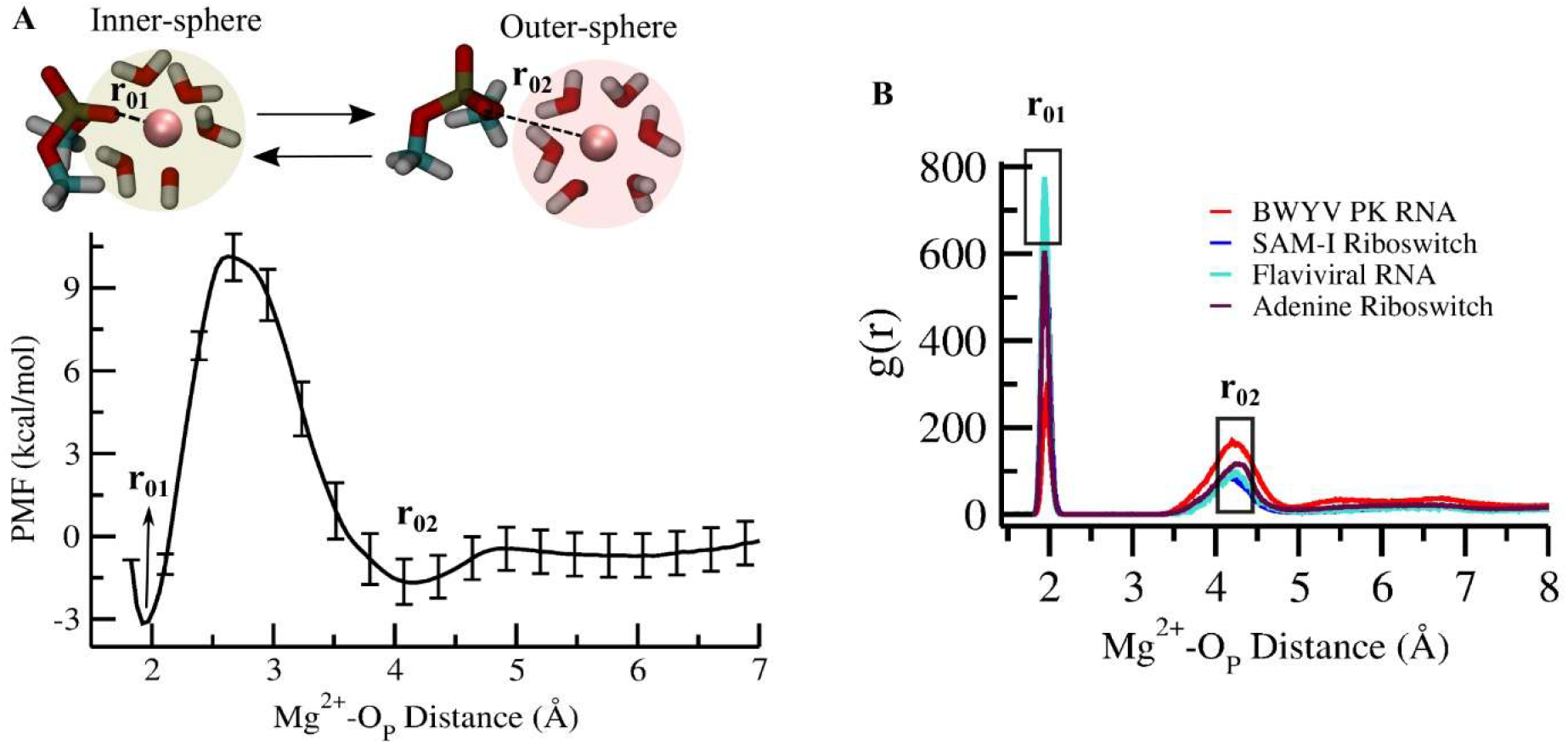
Two complementary approaches to determine the Potential of Mean Force (PMF) and Mg^2+^-RNA coordination modes. (A) PMF profile as a function of the Mg^2+^-O_P_ (oxygen of phosphate) distance, obtained via umbrella sampling for a dimethyl phosphate (DMP) model system. Representative structures correspond to inner-sphere and outer-sphere coordination of Mg^2+^. (B) Radial distribution function (RDF) analysis of Mg^2+^-O_P_ distances in four RNA systems: BWYV pseudoknot, SAM-I riboswitch, flaviviral RNA, and adenine riboswitch, used to identify distinct coordination modes. Both methods consistently identify inner-sphere and outer-sphere distances at r_01_ ≈ 1.9 Å and r_02_ ≈ 4.2 Å, respectively.

To assess whether these features persist in realistic RNA environments, we additionally performed explicit-solvent molecular dynamics simulations of four structurally diverse RNAs: the BWYV pseudoknot (PDB ID: 437D), SAM-I riboswitch (PDB ID: 2GIS), flaviviral RNA (PDB ID: 4PQV), and adenine riboswitch (PDB ID: 1Y26)(10, 44–46). RDFs calculated between Mg^2+^ ions and phosphate oxygens display two characteristic peaks corresponding to distinct coordination geometries, in agreement with crystallographic observations(47) (Figure 1B).

The potential of mean force (PMF) corresponding to a given radial distribution function, g(r), can be obtained via Boltzmann inversion, a widely used approach for deriving coarse-grained interaction potentials(48, 49), as:

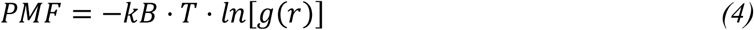

where k_B_ is the Boltzmann constant, and T is the temperature. The PMFs derived from RDF analysis closely mirror those obtained from umbrella sampling (**Figure S1**), providing independent validation of the underlying Mg^2+^-phosphate free-energy landscape. Both approaches consistently identify two characteristic coordination distances: an inner-sphere minimum near 1.9 Å and an outer-sphere minimum near 4.2 Å.

### PMF-Guided Modelling of Mg^2+^ Coordination Exchange

The two PMF minima provide the physical basis for constructing the explicit Mg^2+^ interaction term in STEM. To represent both coordination states and the dynamic exchange between them, we introduce a multi-Gaussian ion-exchange potential,

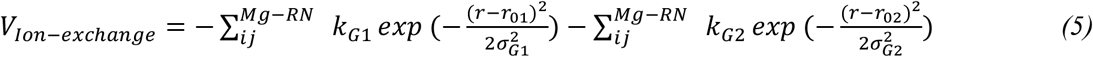

where *r*_*01*_ = 1.9 Å and *r*_*02*_ = 4.2 Å correspond to the experimentally and computationally validated inner-sphere and outer-sphere coordination distances, respectively.

The complete explicit Mg^2+^ interaction potential is then expressed as

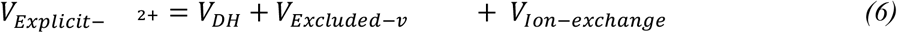

Where, *V*_*DH*_ describes long-range electrostatic interactions through the Debye–Hückel formalism. *V* _*Excluded−v*_ is the excluded volume effect for explicitly treated inner-sphere Mg^2+^ ions. *V*_*Ion−exchange*_ captures transitions between the two coordination states.

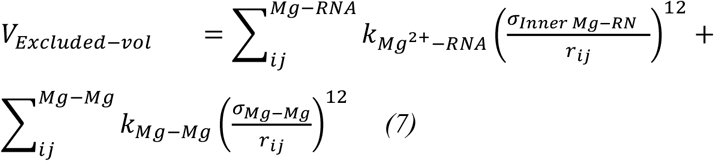

The inner-sphere minimum position set at 1.9 Å, as we defined the excluded volume for inner-sphere Mg^2+^ interactions with RNA (*σ*_*Inner Mg−R*_) as 1.7 Å based on the location of the inner-sphere minimum in the PMF profile. The transition barrier governing coordination-state exchange is controlled by the Gaussian amplitudes, *k*_*G1*_, *k*_*G2*_ and the excluded volume parameter 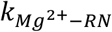. All parameter values are listed in **Table S1**.

### Charge Representation and Initial Validation

To better resolve site-specific Mg^2+^-phosphate interactions, we refined the charge representation used in our previous electrostatic models. Rather than assigning a net charge of (−1) to the phosphate group as a single unit, the charge was distributed equally between the two non-bridging phosphate oxygens (O_P1_ and O_P2_), each carrying a charge of (−0.5). Mg^2+^ ions retained their formal charge of (+2).

As an initial validation, STEM successfully reproduces the characteristic Mg^2+^-phosphate RDF peaks at ∼1.9 Å and ∼4.2 Å corresponding to inner-sphere and outer-sphere coordination states (**Figure S2**) for a number of RNA systems. Direct analysis of Mg^2+^ trajectories further reveals repeated transitions between these states in flaviviral RNA simulations (**Figure S3**), demonstrating that the model captures the intended ion-coordination exchange dynamics.

Subsequent validation against a broader range of experimental observables is presented in the Results section.

## Results

### STEM Captures Both the Thermodynamic and Structural Consequences of RNA–Ion Interactions

Having incorporated dynamic Mg^2+^-coordination exchange into the STEM Hamiltonian, we first examined whether the model quantitatively reproduces experimentally measurable properties of the RNA ion atmosphere. Because Mg^2+^-mediated stabilization arises not only from site-specific interactions but also from the collective accumulation of ions around RNA, an appropriate benchmark is the preferential interaction coefficient, (Γ_2+_), which directly quantifies excess Mg^2+^ associated with an RNA molecule.

The ion atmosphere surrounding RNA contains a population of excess ions whose abundance depends on the bulk ionic conditions while collectively compensating the negative charge of the phosphate backbone. Experimentally, excess Mg^2+^ accumulation has been quantified using fluorescent chelators and equilibrium dialysis measurements, which report the preferential interaction coefficient, Γ_2+_ (3, 18, 50). As a colligative property of the RNA–ion system, Γ_2+_ represents the excess number of Mg^2+^ ions associated with RNA relative to the bulk solution at a given Mg^2+^ concentration (C_2+_). Under conditions where monovalent ions dominate the ionic environment c_+_ ≫ c_2+_, the Mg^2+^-RNA interaction free energy is related to Γ_2+_ through the integral expression: 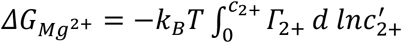, where *k*_*B*_ is the Boltzmann constant, and *T* is the temperature(51). This establishes a direct connection between excess ion accumulation and the underlying electrostatic free energy of RNA–ion association.

We therefore calculated Γ_2+_ from STEM simulations for several structurally distinct RNA systems, including the adenine riboswitch (50 mM KCl; **Figure 2A, B**), the BWYV pseudoknot (54 mM KCl; Figure **2C, D**), and the 58-nt rRNA fragment (150 mM KCl; **Figure 2E, F**). Across all systems, STEM quantitatively reproduces the experimentally measured dependence of Γ_2+_ on Mg^2+^ concentration reported by Draper and co-workers(3, 18, 50). The agreement spans a broad range of RNA architectures and monovalent-salt conditions, demonstrating that the model accurately captures the collective electrostatic properties of the RNA ion atmosphere. Small deviations are observed only at very low Mg^2+^ concentrations (0.1–0.2 mM) for the adenine riboswitch and the 58-nt rRNA fragment. Importantly, STEM faithfully reproduces the experimentally observed behaviour throughout the physiologically relevant concentration regime (0.4–1 mM [Mg^2+^]), where Mg^2+^-dependent RNA folding and stabilization are most pronounced (**Figure 2B, F**). The ability to recover preferential interaction coefficients across multiple RNA systems indicates that STEM accurately captures the free-energy balance governing Mg^2+^ accumulation and electrostatic stabilization around RNA.

**Figure 2:**
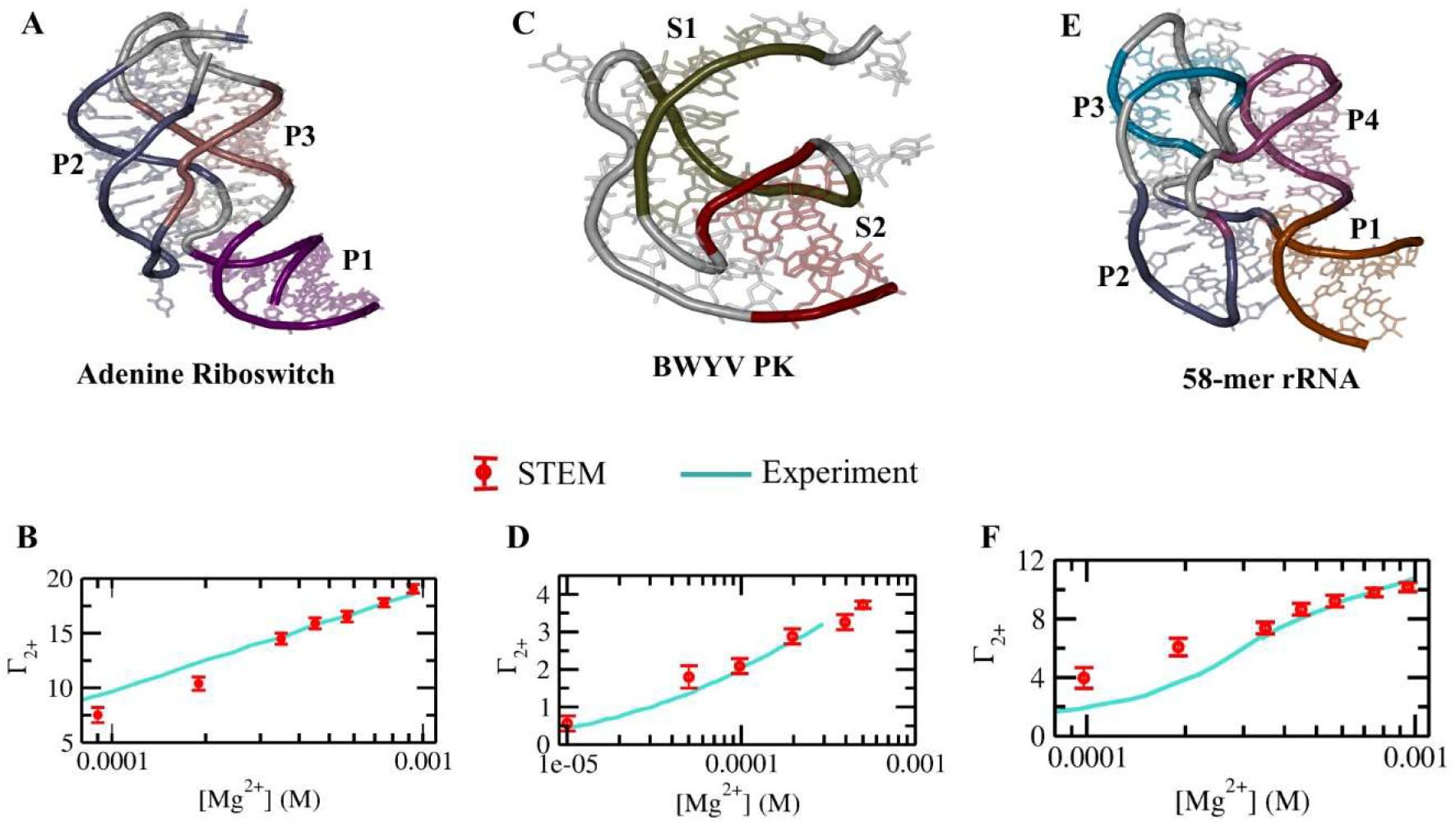
Agreement between STEM simulation and experiments for preferential interaction coefficient (Γ_2+_), estimating excess Mg^2+^ ion accumulation around various RNA structures. (A-F) STEM simulation-predicted excess Mg^2+^ ions over a range of Mg^2+^ concentrations, along with the RNA structures for: (A, B) the adenine riboswitch at 50 mM KCl. (C, D) Beet Western Yellow Virus (BWYV) pseudoknot at 54 mM KCl. (E, F) a 58-nucleotide ribosomal RNA fragment at 150 mM KCl. Experimental excess ion data (by using a fluorescent chelator) are shown as solid lines, and simulation results as dots with error bars.

While preferential interaction coefficients probe the thermodynamics of the ion atmosphere, they do not directly assess its structural consequences. We therefore next examined whether STEM reproduces experimentally observed Mg^2+-^induced RNA compaction. The adenine riboswitch provides an ideal test system because its solution structure has been extensively characterized by small-angle X-ray scattering (SAXS) under both Mg^2+^-free and Mg^2+^-containing conditions(3). We performed equilibrium STEM simulations at 50 mM KCl in the presence and absence of Mg^2+^ and monitored the resulting distributions of the radius of gyration (R_g_) (**Figure S4A**). Experimentally, the riboswitch adopts a compact native-like conformation with a radius of gyration (R_g_) of 20.3 Å in the presence of Mg^2+^, whereas removal of Mg^2+^ produces a substantially expanded ensemble with an R_g_ of 24.9 Å (45). STEM faithfully reproduces this transition, yielding mean (R_g_) values of 20.3 Å at 1 mM Mg^2+^ and 24.4 Å in the absence of Mg^2+^ (**Figure S4**). Thus, the model not only captures the thermodynamics of excess ion accumulation but also the resulting structural reorganization of RNA in solution.

Taken together, these results demonstrate that STEM accurately describes both the collective properties of the RNA ion atmosphere and its structural consequences, providing a robust foundation for investigating how Mg^2+^-coordination exchange influences RNA folding pathways and conformational dynamics.

### STEM Identifies Site-specific Mg^2+^ Binding Sites in RNA

Having established that STEM accurately reproduces both the thermodynamic and structural signatures of the RNA ion atmosphere, we next asked whether the model can resolve the site-specific Mg^2+^ interactions that stabilize RNA tertiary structure. Crystallographic studies have shown that a small number of inner-sphere Mg^2+^ ions often occupy highly specific locations within folded RNAs, where they stabilize tertiary contacts and, in some cases, directly support biological function(10, 52). Accurately identifying these binding sites therefore provides a stringent test of the model’s ability to capture local ion-mediated interactions beyond collective electrostatic effects. Because STEM explicitly incorporates inner-sphere Mg^2+^ coordination, it enables direct prediction of Mg^2+^-binding hotspots from RNA structure and dynamics. To assess its predictive capability, we examined three RNAs for which crystallographically resolved inner-sphere Mg^2+^ binding sites are available: the BWYV pseudoknot (PK), a 58-nt rRNA fragment, and a flaviviral RNA(53–55).

To eliminate potential bias from experimentally resolved ions, all crystallographic Mg^2+^-mediated contacts were removed prior to simulation. Mg^2+^ ions were then randomly distributed within the simulation box, and equilibrium simulations were performed using the full STEM Hamiltonian. Site-specific Mg^2+^-binding propensities were quantified from residue-resolved inner-sphere coordination frequencies between Mg^2+^ ions and phosphate oxygens (O_P_). Details of the analysis are provided in the Supporting Information.

Figure 3 compares the binding-site probabilities predicted by STEM with experimentally resolved Mg^2+^ locations. Residues exhibiting high coordination frequencies > 0.6)) were classified as probable inner-sphere Mg^2+^-binding sites. Remarkably, the simulations recover the principal crystallographic binding sites in all three RNA systems. For the BWYV pseudoknot, the crystal structure contains a chelated Mg^2+^ ion positioned near the GTP-binding region involving residues GTP1 and G2 (47). STEM predicts a pronounced binding hotspot at the same location (Figure 3A, B). Similarly, for the 58-nt rRNA fragment, the experimentally resolved Mg^2+^ coordinated near residues C21, A22, and U43 (48) is accurately reproduced by the simulations (**Figure 3C, D**). In addition to this known site, STEM identifies a second high-probability binding region near G11, suggesting the existence of a transient Mg^2+^-stabilized interaction that may influence RNA conformational dynamics. Finally, for the flaviviral RNA, the crystallographic Mg^2+^ site located between residues C5 and C23 (55) is recapitulated by a strongly populated binding region involving residues C5, C6, and A24 in the simulations (**Figure 3E, F**).

**Figure 3:**
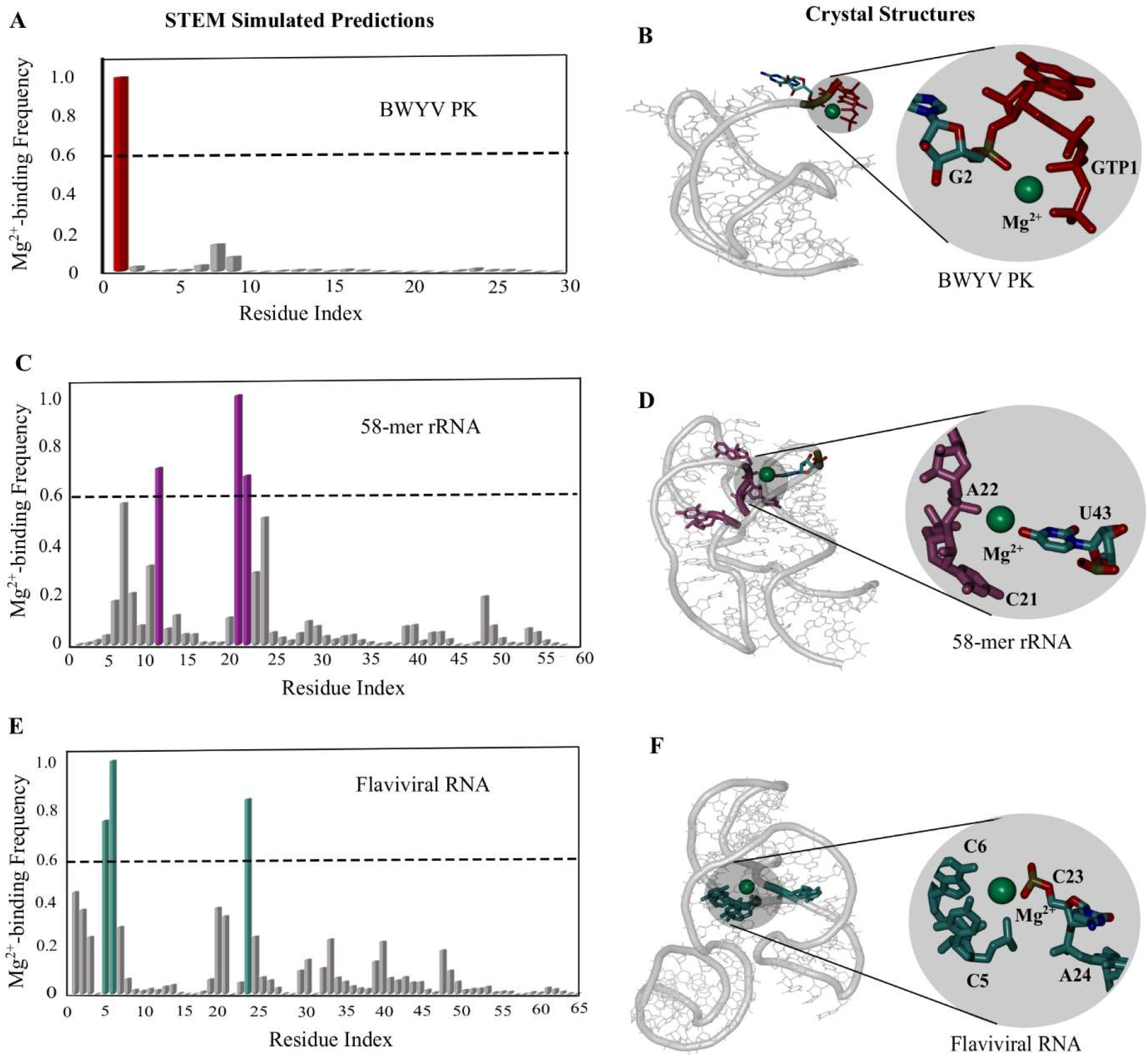
Prediction of Mg^2+^ binding sites across various RNAs by STEM simulation. (A) Residue-wise Mg^2+^ binding frequency for the BWYV pseudoknot (PK). (B) Crystal structure of the BWYV PK highlighting chelated Mg^2+^ ions. (C) Residue-wise Mg^2+^ binding frequency for the 58-mer rRNA fragment. (D) Corresponding crystal structure of 58-mer rRNA with chelated Mg^2+^ ions. (E) Residue-wise Mg^2+^ binding frequency for the flaviviral RNA. (F) Corresponding crystal structure of flaviviral RNA with chelated Mg^2+^ ions. In each structural representation, residues with predicted binding frequency > 0.6 are highlighted with the same colour combination of the bar plot, and experimentally resolved Mg^2+^ ions are shown as green spheres.

The ability of STEM to recover experimentally observed inner-sphere Mg^2+^ binding sites without imposing any prior knowledge of their locations demonstrates that the model captures the physical determinants governing site-specific ion recognition. Moreover, the emergence of additional high-probability Mg^2+^-binding regions suggests that the simulations may reveal transient or dynamically populated coordination sites that are difficult to resolve in static crystallographic structures. These sites become particularly relevant in the context of RNA conformational fluctuations and folding transitions, which we investigate in the following sections.

### STEM Captures Ion Chelation Induced Structural Intermediates Exploring RNA Folding Energy Landscape

So far, we have validated STEM by accurately capturing RNA structural features, the surrounding ion atmosphere, and inner-sphere Mg^2+^ binding site predictions across various RNAs. We next investigate the functional role of inner-sphere Mg^2+^ ions in the conformational transitions of RNA, with particular emphasis on exploring intermediate states relevant to RNA folding. To this end, we focus on 58-mer rRNA, motivated by an earlier experimental study from Draper and colleagues, which showed that an important intermediate (I) state of this RNA lacks inner-sphere Mg^2+^ coordination, while the transition to the native (N) folded state involves the appearance of a single chelated Mg^2+^ ion(18).

To explore this mechanism at atomistic detail, we evaluated the folding free energy landscape by performing umbrella sampling simulations(39). A series of 28 overlapping windows was employed to ensure adequate sampling across conformational space, with histograms constructed to confirm sufficient overlap. Details of the simulation setup and free energy calculations are provided in the Supporting Information. The global fraction of native contacts (Q_Global_) was used as the order parameter to characterize the folding process under physiological ionic conditions (100 mM KCl and 2 mM Mg^2+^), as shown in **Figure 4A**. To isolate the specific role of inner-sphere Mg^2+^, we removed all Mg^2+^-mediated contacts from the crystal structure before simulation to eliminate any bias from pre-formed binding motifs.

**Figure 4:**
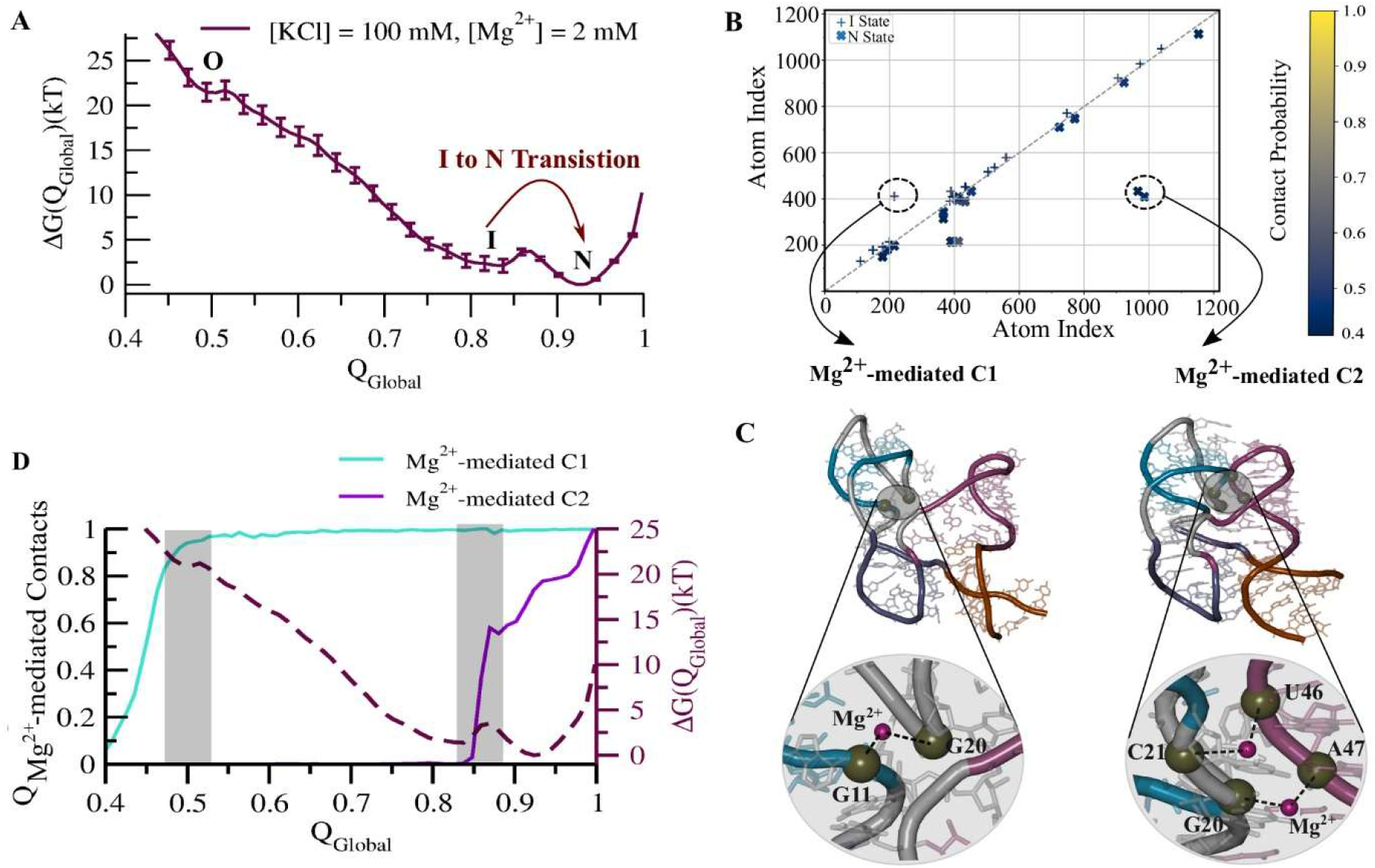
Folding free-energy landscape and inner-sphere Mg^2+^-mediated contact probability maps for the Intermediate (I) and Native-like (N) states of the 58-mer rRNA, illustrating ion-mediated stabilization along the folding pathway. (A) Folding free energy profile as a function of the global fraction of native contacts (Q_Global_), revealing three barrier-separated minima corresponding to distinct conformational states. (B) Inner-sphere Mg^2+^-mediated contacts between non-local RNA atoms in the intermediate (I) and native-like (N) states. The upper diagonal of the contact map corresponds to the I state, and the lower diagonal to the N state. (C, D) Structural representation of the I state, highlighting non-local interactions bridged by inner-sphere Mg^2+^ ions (Mg2+-mediated C1) and the structural representation of the N state, showing two regions where inner-sphere Mg^2+^ ions mediate non-local contacts (Mg2+-mediated C2). (D) Sequential folding of Mg^2+^-mediated contacts, shown as a function of Q_Global_. Here, the highlighted transparent layers indicate significant barrier-crossing events (O to I and I to N).

The resulting free energy profile revealed three well-defined minima, each separated by energy barriers, corresponding to distinct conformational ensembles: (i) the fully folded native-like state (N), (ii) an intermediate state (I), and (iii) a partially unfolded or open state (O) (**Figure 4A**). Structurally, the O state corresponds to a conformation where the P3 helix is only partially formed; the I state has all secondary structures intact, but lacks tertiary interactions at the P1– P2 and P3–P4 junctions; the N state represents a compact, native-like conformation stabilized by both secondary and tertiary interactions (**Figure S5**). Importantly, these conformational states and their ordering along the folding landscape are consistent with experimental observations reported by Draper and co-workers (4). To compare the structural properties of the folded native-like (N) and intermediate (I) states, we calculated the radius of gyration (Rg), which shows good agreement with the small-angle X-ray scattering (SAXS) measurements reported by Draper and co-workers (**Figure S6**).

To elucidate the molecular basis of the I → N transition, we computed inner-sphere Mg^2+^-mediated contact probability maps for both the I and N states. The details of the inner-sphere Mg^2+^-mediated contact probability maps are described in our recent work(45). In the resulting contact map (**Figure 4B**), the upper triangle represents Mg^2+^-mediated contacts in the I state, whereas the lower triangle corresponds to those in the N state. Details of the contact analysis are provided in the Methods section.

This analysis revealed two distinct Mg^2+^-coordinated interactions: (i) Mg^2+^-mediated Contact 1 (C1), which is present in both the I and N states and involves coordination between residues G11 and G20 (**Figure 4C**); (ii) Mg^2+^-mediated Contact 2 (C2), which is unique to the N state and arises from inner-sphere Mg^2+^ coordination bridging residues C21–U46 and G20–A47 (**Figure 4C**).

To connect these interactions to folding dynamics, we analyzed the free-energy landscape in conjunction with the formation of these specific Mg^2+^-mediated contacts. This analysis reveals two distinct barrier-crossing events along the folding pathway, corresponding to the emergence of the Mg^2+^-mediated contacts C1 and C2. These transitions occur at Q_Global_ ≈ 0.5 and Q_Global_ ≈ 0.85, respectively (**Figure 4D**). The formation of Mg^2+^-mediated C1 marks the initiation of tertiary folding, whereas the Mg^2+^-mediated C2 accompanies the completion of native-like tertiary folding. The exclusive presence of C2 in the N state suggests its critical role in stabilizing long-range tertiary interactions during the I → N transition. Together, these results provide mechanistic insight into how inner-sphere (chelated) Mg^2+^ ions facilitate the final stages of RNA folding by mediating long-range tertiary interactions.

In addition to characterizing these barrier-crossing events associated with tertiary contact formation, we examined the hierarchical folding pathway of individual structural elements of RNA. To this end, we calculated the fraction of native intra-segment contacts (Q_Regional_) for each helical segment as a function of the global native contacts (Q_Global_) along the free energy landscape. The results reveal a defined sequence of folding events: folding initiates at helix P3, followed by P2, then P4, and finally P1 (**Figure S5**). Interestingly, during the transition from the open (O) to intermediate (I) state, helix P4 exhibits transient backtracking near Q_Global_ ≈ 0.61. This partial unfolding appears to facilitate the correct formation of P2, indicating a cooperative interplay between helices during intermediate folding. The final stabilization of P1 marks the completion of folding and the attainment of the native-like state.

### STEM Reveals Dynamic Ion-Coordination-Exchange Controlling RNA’s Conformational Dynamics

Our free energy analysis indicates that inner-sphere Mg^2+^-mediated interactions play a crucial role in the transition from the intermediate (I) state to the native-like (N) state. These results point to a dynamic equilibrium, or conformational breathing, between the I and N states. Building on experimental observations by Draper and co-workers (18), these findings suggest that the transition is regulated by the exchange of Mg^2+^ ions between outer-sphere and inner-sphere coordination states, a mechanism that is explicitly encoded in the STEM Hamiltonian.

To test this hypothesis, we performed multiple unbiased equilibrium simulations of the 58-mer rRNA at physiological ionic conditions (100 mM KCl and 2 mM Mg^2+^). Across all simulation trajectories, the model captured spontaneous transitions between the I and N states, revealing their intrinsic breathing behaviour. A representative trajectory illustrating these transitions is provided in **Movie S1**. Structural analysis identified two key segments—P3* (residues G17– G23) proximal to helix P3 and P4* (residues C41–G48) near helix P4—that distinguish the I and N conformations. The distribution of centre-of-mass (COM) distance between P3* and P4* exhibits pronounced fluctuations, directly reporting on the breathing motion between these two states (**Figure 5A**).

**Figure 5:**
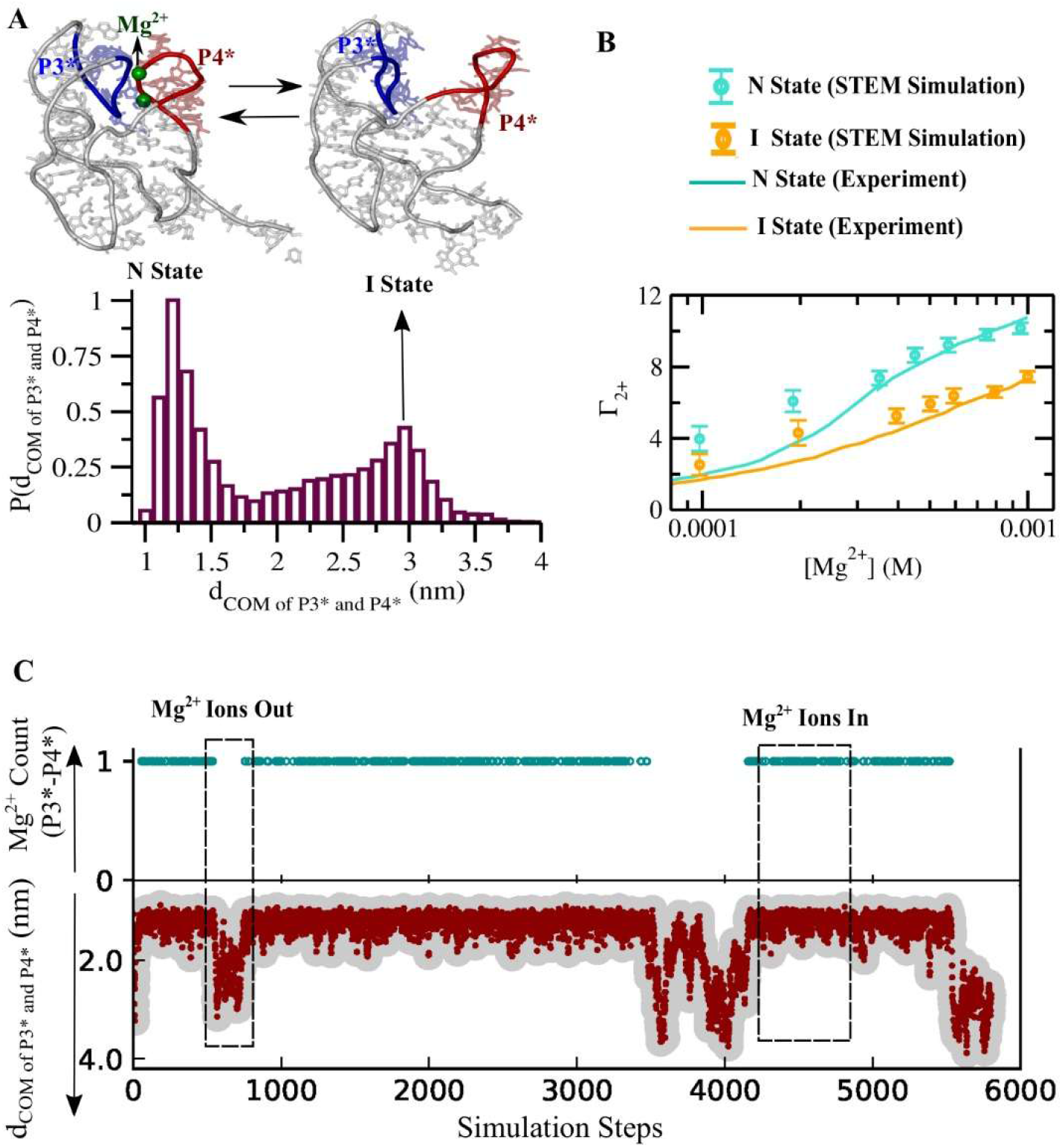
Mg^2+^ ion exchange modulates conformational dynamics of the 58-mer rRNA fragment. (A) Distribution of centre-of-mass (COM) distances between the P3* and P4* segments, calculated from multiple simulation trajectories. Representative structural snapshots depict the intermediate (I) and native-like folded (N) states, as previously identified in experiments. (B) Predicted excess Mg^2+^ ions across a range of Mg^2+^ concentrations for the I and N states, revealing distinct ion accumulation patterns associated with each conformation. (C) Time evolution of COM distances between P3* and P4*, illustrating a breathing-like transition from open to closed conformations, modulated by Mg^2+^ ion exchange. In the ion exchange calculation, a value of 0 indicates the absence of inner-sphere Mg^2+^, while a value of 1 denotes its presence.

To assess the physiological relevance of these dynamics, we computed the preferential interaction coefficient (Γ_2+_) for both the I and N states at 150 mM KCl over a range of Mg^2+^ concentrations and compared the results with experimental measurements from Draper and co-workers (4). The simulations reproduce the experimental trends, demonstrating that the I state is associated with fewer excess Mg^2+^ ions than the N state (**Figure 5B**), consistent with the absence of stable inner-sphere coordination in the intermediate ensemble.

From our earlier analysis of inner-sphere Mg^2+^-mediated contacts, we identified Mg^2+^-mediated Contact 2 (C2)—formed by interactions between C21–U46 and G20–A47—as a stabilizing interaction unique to the N state (**Figure 4B, C**). To directly link ion exchange at this site to conformational breathing, we monitored the occupancy of inner-sphere Mg^2+^ at C2 across equilibrium trajectories, assigning values of 1 (bound) and 0 (unbound). When plotted against the P3*–P4* COM distance (**Figure 5C**), ion occupancy exhibits a clear correlation with conformational state: transitions between open and closed configurations are consistently accompanied by the binding or release of Mg^2+^ at C2.

Together, these results establish a direct mechanistic coupling between inner-sphere Mg^2+^ coordination exchange and RNA conformational dynamics. The correlation between breathing motion and Mg^2+^ coordination at C2 provides strong evidence that Mg^2+^ coordination exchange plays an essential role in RNA folding transitions. Overall, the STEM Hamiltonian—by explicitly allowing dynamic exchange between outer-sphere and inner-sphere coordination states—captures the microscopic basis of ion-driven conformational transitions in RNA.

## Discussion

The Structural Topology-based Electrostatic Model (STEM) was developed to address a longstanding challenge in RNA electrostatics: how to describe, within a computationally tractable framework, the dynamic exchange between distinct ion-coordination states and its coupling to RNA conformational transitions. By combining explicit Mg^2+^ ions with an implicit treatment of monovalent ions through Generalized Manning Counterion Condensation theory(33), STEM captures the essential physics of ion condensation and site-specific divalent-ion coordination while remaining efficient enough to access large-scale folding dynamics. Incorporation of a PMF-derived ion-exchange potential further enables the model to directly connect Mg^2+^ coordination to RNA folding dynamics.

Across multiple RNA systems, STEM quantitatively reproduces experimental observables spanning different scales of RNA–ion interactions, including preferential Mg^2+^ accumulation, Mg^2+^-dependent RNA compaction, and site-specific inner-sphere Mg^2+^ binding patterns. More importantly, application of STEM to the 58-nt rRNA fragment reveals a mechanistic role for Mg^2+^ coordination exchange during folding. Consistent with Draper and coworkers’ experiments(18, 57), the transition from the intermediate to the native state is associated with the emergence of a specific inner-sphere Mg^2+^-mediated tertiary contact. STEM further shows that reversible formation and disruption of this contact give rise to spontaneous conformational breathing between the intermediate and native ensembles. Together, these results support a view in which Mg^2+^ ions are not merely passive electrostatic stabilizers, but active regulators of RNA conformational transitions.

This picture extends the conventional view of the RNA ion atmosphere as a static electrostatic background. Instead, our results suggest that RNA conformation and ion organization are dynamically coupled, such that transitions between outer-sphere and inner-sphere Mg^2+^ coordination can reshape the folding free-energy landscape. The importance of distinguishing these coordination modes has been recognized in earlier theoretical and computational studies(13, 14, 58), including PMF-based coarse-grained models that incorporate both inner- and outer-sphere divalent-ion interactions with phosphate groups(29, 31). However, those studies primarily focused on equilibrium ion distributions and folding thermodynamics rather than on whether exchange between coordination states is itself coupled to conformational transitions. In this sense, the central advance of STEM is not simply the inclusion of multiple coordination modes, but the demonstration that their exchange can be mechanistically linked to RNA folding-state interconversion and conformational breathing.

The ability to explicitly model ion-coordination exchange also helps address a gap in current RNA structure-prediction strategies. Recent computational methods have markedly improved the prediction of RNA contacts and native folds(59–61), but they generally lack an explicit description of how ions stabilize and modulate RNA conformations. STEM provides a complementary physics-based framework that can identify likely ion-binding sites, describe both collective and site-specific ion-mediated interactions, and resolve folding free-energy landscapes and competing conformational ensembles beyond the native basin.

Several limitations of the present framework should be noted. STEM has been developed primarily for Mg^2+^-mediated RNA folding and does not yet explicitly account for competition among multiple divalent ion species. In addition, although the PMF-based ion-exchange potential captures transitions between inner-sphere and outer-sphere coordination, solvent remains implicit and therefore cannot resolve detailed water-mediated rearrangements during ion binding and release. The current implementation is also focused on isolated RNAs and does not yet include protein partners, metabolites, or macromolecular crowding effects. Despite these limitations, STEM establishes a mechanistic and predictive framework for investigating ion-dependent RNA folding by unifying RNA topology, electrostatics, and dynamic Mg^2+^ coordination exchange within a single computational model.

## Conclusion

In conclusion, we developed the Structural Topology-based Electrostatic Model (STEM), a hybrid framework that integrates RNA structural topology, implicit monovalent-ion screening, and explicit Mg^2+^ coordination within a single computationally efficient model. By incorporating dynamic exchange between outer-sphere and inner-sphere Mg^2+^ coordination states, STEM moves beyond static descriptions of RNA electrostatics and enables direct investigation of how ion-mediated interactions shape RNA conformational landscapes. Across multiple RNA systems, STEM accurately reproduces experimental signatures of the RNA ion atmosphere, including preferential Mg^2+^ accumulation, Mg^2+^-dependent RNA compaction, and site-specific inner-sphere Mg^2+^ binding. Applied to the 58-nt rRNA fragment, the model further reveals that the transition from the intermediate to the native state is coupled to the emergence of a specific chelated Mg^2+^-mediated tertiary contact and to dynamic conformational breathing driven by Mg^2+^ coordination exchange. Together, these results support a view of RNA folding in which ions are not merely passive electrostatic stabilizers but active participants in sculpting the folding free-energy landscape. By linking ion-coordination dynamics to RNA conformational transitions, STEM provides a mechanistic framework for predicting dynamic RNA ensembles and their ion-dependent folding pathways, with broader implications for understanding ribozymes, riboswitches, and other functional RNAs under physiological ionic conditions.

## ACKNOWLEDGEMENTS

S.R., as an academic visiting faculty member of The Center for Theoretical Biological Physics (CTBP), Rice University, particularly thanks Rice Supercomputing facility for its computational support. S.R. also thanks Dr. Ryan Hayes for providing the source code for the previous model, and Prof. Paul Whitford and Dr. Heiko Lammert for several insightful discussions. This work was supported by Anusandhan National Research Foundation (Grant no: ANRF/ARG/2025/008640/CS) and partly by the Department of Biotechnology (DBT) (Grant No. and BT/PR40192/BTIS/137/69/2023), Govt. of India. J.N.O. and K.Y.S. acknowledge NIH NIGMS grant R01GM110310. A. M. acknowledges support from the CSIR-NET fellowship.

## Supporting Information for

### Electrostatic Effect as Potential Function for Implicit KCl (*V*_*Implicit*−*KCl*_)

The implicit effect of KCl has been incorporated into the Hamiltonian through the term *V*_*Implicit*−*KCl*_. Monovalent K^+^ ions are modelled implicitly using GMCC theory(1), allowing for computational efficiency. Following the GMCC framework, the local charge density of implicit K^+^ ions is represented as a smeared Gaussian shell around the centre of mass of each negatively charged phosphate group. This Gaussian shell defines an effective volume that excludes the phosphate’s steric core and represents the region of condensed implicit ions. For the implicit treatment of KCl, the model accounts for three types of electrostatic interactions: point charge–point charge, point charge–Gaussian, and Gaussian–Gaussian. Each of these interactions is treated in terms of the Debye–Hückel potential, and together they contribute to the total electrostatic energy of the system as,

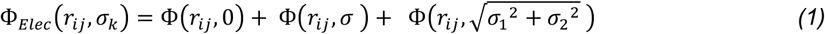

Here, the overall electrostatic free energy 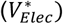 governing all the electrostatic interactions Φ_*Elec*_ (*r*_*ij*_, *σ*_*k*_) will compete with mixing free energy 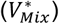.

Competition between electrostatic free energy and mixing free energy identifies the counter-ion condensation on negatively charged phosphate, similar to classical Manning theory(2). In our STEM, to maintain the intermediate KCl salt concentration (∼100 mM), we have added the effect of screening ion density with implicit condensed ions.

The density of condensed ions of type ‘s’, around each negatively charged phosphate, ‘i’ is modelled as a sum of two normalized Gaussian distributions to get charge heterogeneity around each phosphate group of RNA. Therefore, the electrostatic free energy term appears as,

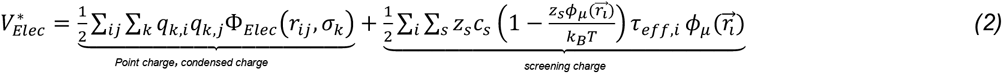

The indices, a and b define the pair-wise Debye-Hückel interactions (*ϕ*) between any of the charge types. The distance between particles i and j is *r*_*ij*_. *z*_s_ is the charge number and *c*_s_ is the concentration of a specific ion, s. For particle i, the condensed charges are, *q*_*µ,i*_ = ∑_*s*_ *z*_*s*_*μ*_*i,s*_ and *q*_*η,i*,_ = ∑_*k*_ *z*_*s*_*η*_*i,s*_. We reckoned with an effective Gaussian shell volume such as, *τ*_µ,*i*_ − *τ*_*η,i*,_ = *τ*_eff, *i*_.

The mixing free energy within the effective Gaussian shell volume is given as,

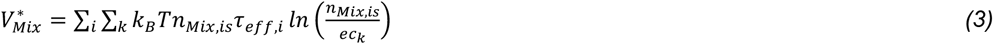

where e is the Euler’s number and *n*_*Mix,is*_ is the local mixing density.

### STEM Simulation Details

Atomic coordinates and condensation variables (*μ*_*i*,s_ and *η*_*i*,s_) are evolved with Langevin dynamics with a time step of 0.001*τ*_*R*_. We used an underdamped condition for rapid sampling. For explicit particles, reduced mass of 1*μ*_*R*_ and drag coefficient of 1 *τ* ^−1^ are used. Condensation parameters, *μ*_*i*,k_ and *η*_*i*,k_ are given a mass of 15 *μ*_*R*_ nm^2^ and a drag coefficient of 0.05 *τ* ^−1^ nm^2^. Temperature was chosen to 87*T*_*R*_ to capture the breathing dynamics. To ready up different Mg^2+^ composition, we created a large cubic box of length 75 nm. The number of Mg^2+^ molecule included in that box determines the overall concentration of the corresponding solutes. Periodic boundary conditions were applied. Each simulation was run with 200 million of time steps.

The related parameter set and its calibration are available in **Table S1**.

▪ **Values of Parameter Set for the STEM are Given in Table S1**

| Reduced parameter set |  |
| --- | --- |
| $\tau_R$ | 2 ps |
| $\mu_R$ | 15 amu |
| $\epsilon_R$ | 3.75 KJ/mol |
| $T_R$ | 3.75 K |
| Excluded volume parameter |  |
| $\sigma_{inner-Mg-RNA}$ | 0.17 nm |
| $\sigma_{Mg-Mg}$ | 0.56 nm |
| Ion exchange parameter |  |
| $r_{01}$ | 0.19 nm |
| $r_{02}$ | 0.42 nm |
| $k_{G1}$ | $0.3\epsilon_R$ |
| $k_{G2}$ | $0.2\epsilon_R$ |
| $\sigma_{G1}$ | 0.05 |
| $\sigma_{G2}$ | 0.05 |

### Free Energy Calculation and Umbrella Sampling Details

To sample the whole conformation landscape of RNA, we calculated folding free energy using the umbrella sampling method(3), where the fraction of the global native contacts (Q_Global_) is selected as an order parameter. A collection of initial structures is generated for each window of Q_Global_ by adopting a slow pull criterion, ensuring an ion-equilibrated initial structure. However, we again reinitialized the explicit Mg^2+^ ions distribution at each umbrella window to extend the equilibration and ran for 200 million steps. To confirm the substantial overlap in the conformational space, we have used a total of 28 windows along the Q_Global_. Next, the weighted histogram analysis method (WHAM)(4) is used to calculate the thermodynamic free energy landscape, G(Q_Global_).

### Details of the Explicit Solvent Equilibrium Simulation Method

We have created a model system that resemble the RNA backbone to study magnesium-phosphate interaction and exchange phenomena between inner-sphere mono-phosphate coordinated and solvent separated outer-sphere Mg^2+^ ions. Avoiding the topology-specific structural complexity, the inner-sphere mono-phosphate coordinated Mg^2+^ coordination has been studied by taking Mg^2+^-coordinating with a single dimethyl phosphate (DMP) anion.

The initial structures of these Mg^2+^-phosphate complexes were taken from the crystal structure of the SAM-I riboswitch (PDB ID: 2gis)(5). The Avogadro program(6) was used to add hydrogen atoms to make valency adjustments. AMBER(7), a commonly used biomolecular force field, was employed to parameterize for the DMP systems. Force field parameters for AMBER were obtained from the Generalized Amber Force Field (GAFF)(8). The DMP systems were confined to cubic boxes of dimension 5 Å × 5 Å × 5 Å. An optimized TIP3P water model was employed(9), consistent with previous study demonstrating minimal impact of water model variations on Mg^2+^-phosphate kinetic descriptions(10). To ensure system neutrality, appropriate counter-ions were added. Mg^2+^-phosphate interactions have been investigated in a physiologically relevant salt concentration of 100 mM [KCl].

On the other hand, to simulate four RNA systems—the BWYV pseudoknot (PDB ID: 437D), SAM-I riboswitch (PDB ID: 2GIS), flaviviral RNA (PDB ID: 4PQV), and adenine riboswitch (PDB ID: 1Y26)— employing a rigorous equilibration protocol(11–14). The Amber 99 force field(7) with parmbsc0(15) and chiOL3 extensions(16), was employed to generate the atomistic topologies, for all the four RNAs. The systems include inner-sphere/chelated Mg^2+^ ions. All the four RNAs were placed in a cubic simulation box with dimensions of 100 Å × 100 Å × 100 Å. Following DMP systems, the TIP3P water model was used as a solvent model. To neutralize the overall system charge, a suitable number of Mg^2+^, K^+^, and Cl^−^ ions were added, maintaining concentrations of 2 mM [Mg^2+^] and approximately 100 mM [K^+^], thereby simulating a mixed salt ionic environment within the 100 Å × 100 Å × 100 Å box.

#### RNA Equilibration Method

The accurate equilibration of ion distributions around RNA, specifically the outer-sphere and inner-sphere ions, in simulations involving a mixture of counter-ions (K^+^, Mg^2+^) is challenging. This complexity arises from two primary issues: (i) The RNA backbone’s strong electrostatic attraction to Mg^2+^ ions causes rapid condensation of Mg^2+^ onto the RNA without the formation of a proper hydration shell; (ii) Once Mg^2+^ ions interact with the RNA, they may adhere to negatively charged sites, with the energy barrier preventing easy unbinding, even if hydrated. The ions in our simulations are a combination of excess ions that balance the RNA charge and bulk ions, accounting for the physiological ion concentration range. To avoid premature condensation of Mg^2+^ ions onto the RNA, the ions were randomly placed inside the simulation box with larger van der Waals radii. Stochastic dynamics were employed for a 10 ns equilibration, with the RNA constrained and a dielectric constant of 80 to mimic water, ensuring convergence of the electrostatic energy. After filling the simulation box with water, we minimized the system’s energy using the steepest descent method(17) and annealed it to 300 K over 500 ps with positional restraints of 1000 kJ/mol/nm^2^ applied to both Mg^2+^ ions and RNA. Following our early protocol, Mg^2+^ ions were first released and equilibrated for 2 ns, then RNA restraints were progressively reduced from 1000 to 100 to 0 kJ/mol/nm^2^ at constant volume, leading to a total 10 ns NVT equilibration. An additional 10 ns of unrestrained equilibration under constant pressure was performed. After these steps, the systems were subjected to a production MD run.

### Calculation of Binding Sites prediction

The Mg^2+^-binding frequency quantifies how often inner-sphere Mg^2+^ ions approach a partially negatively charged oxygen of phosphate atoms. A specified cut-off distance determines the proximity between a phosphate oxygen and an explicit Mg^2+^ ion. The cut-off distance is determined based on the radial distribution function (RDF) of inner-sphere Mg^2+^ ions around the phosphate oxygen atoms (**Figure S2**).

### Figures

**Figure S1:**
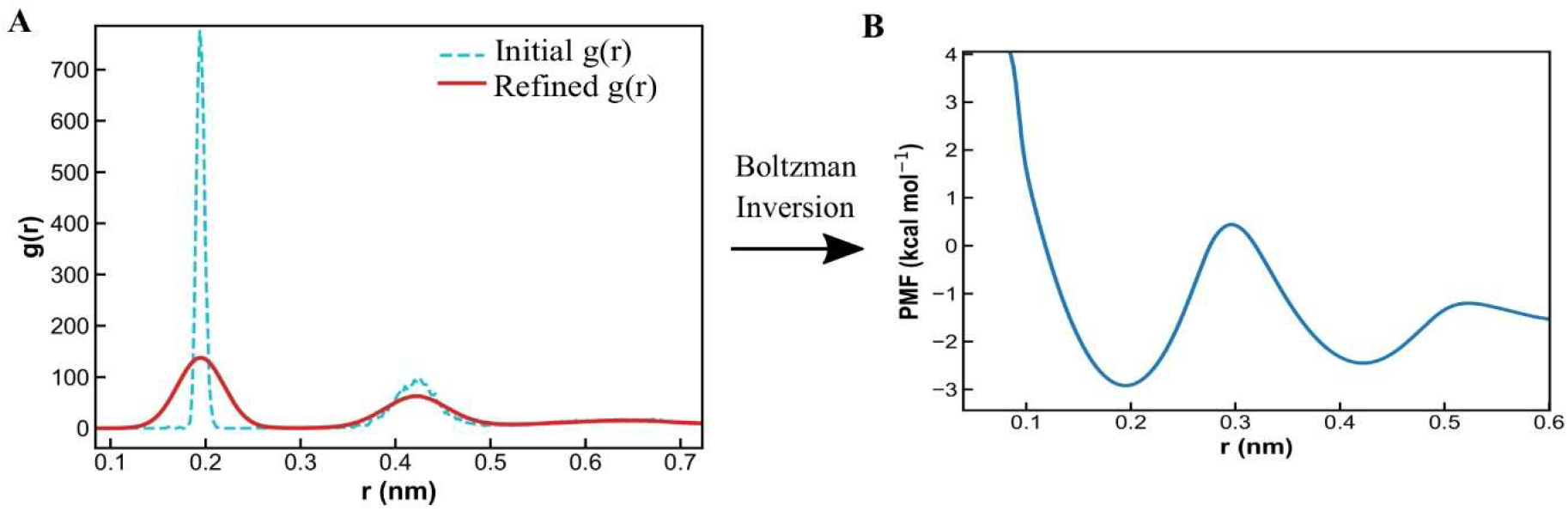
PMF profile obtained using the Boltzmann inversion method. (A) Radial distribution function, g(r), of Mg^2+^–O_P_ distances calculated from explicit-solvent simulations of a flaviviral RNA pseudoknot. The cyan dashed line represents the initial g(r), while the red solid line corresponds to the refined g(r) used for PMF construction. (B) Potential of mean force (PMF) derived from Boltzmann inversion of the refined g(r), revealing distinct inner-sphere and outer-sphere coordination minima at 1.9 Å and 4.2 Å, respectively.

**Figure S2:**
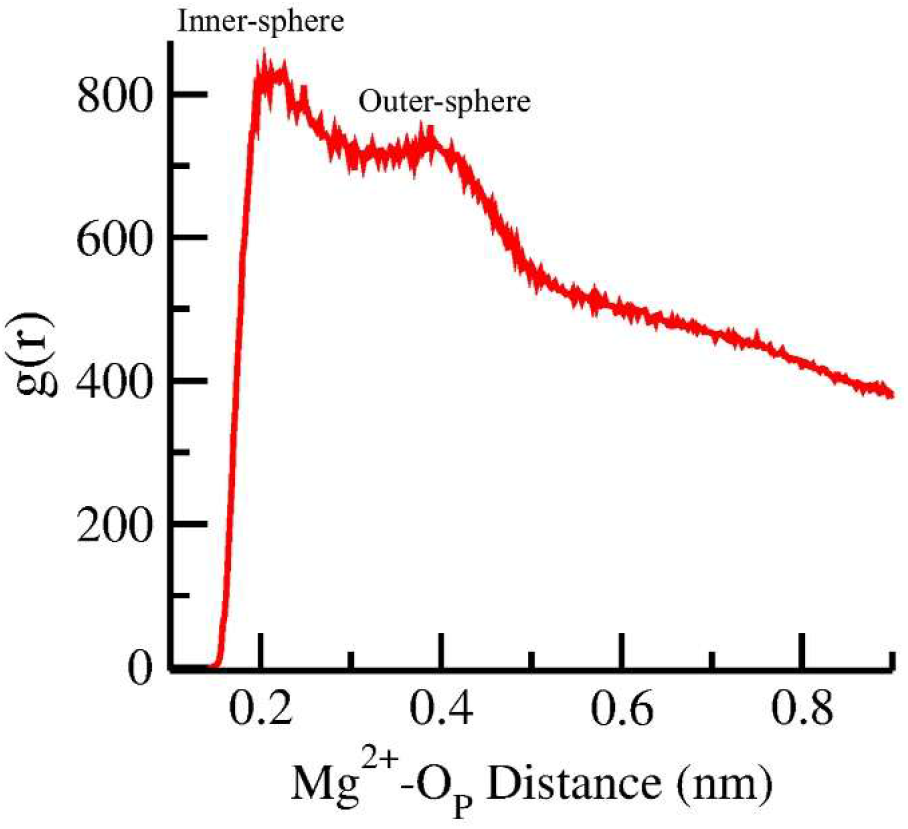
Radial distribution function (RDF) obtained from the STEM framework after incorporating dynamic ion-exchange effects into the Hamiltonian, reproducing the characteristic peaks at ∼1.9 Å and ∼4.2 Å corresponding to inner-sphere and outer-sphere Mg^2+^ coordination, respectively.

**Figure S3:**
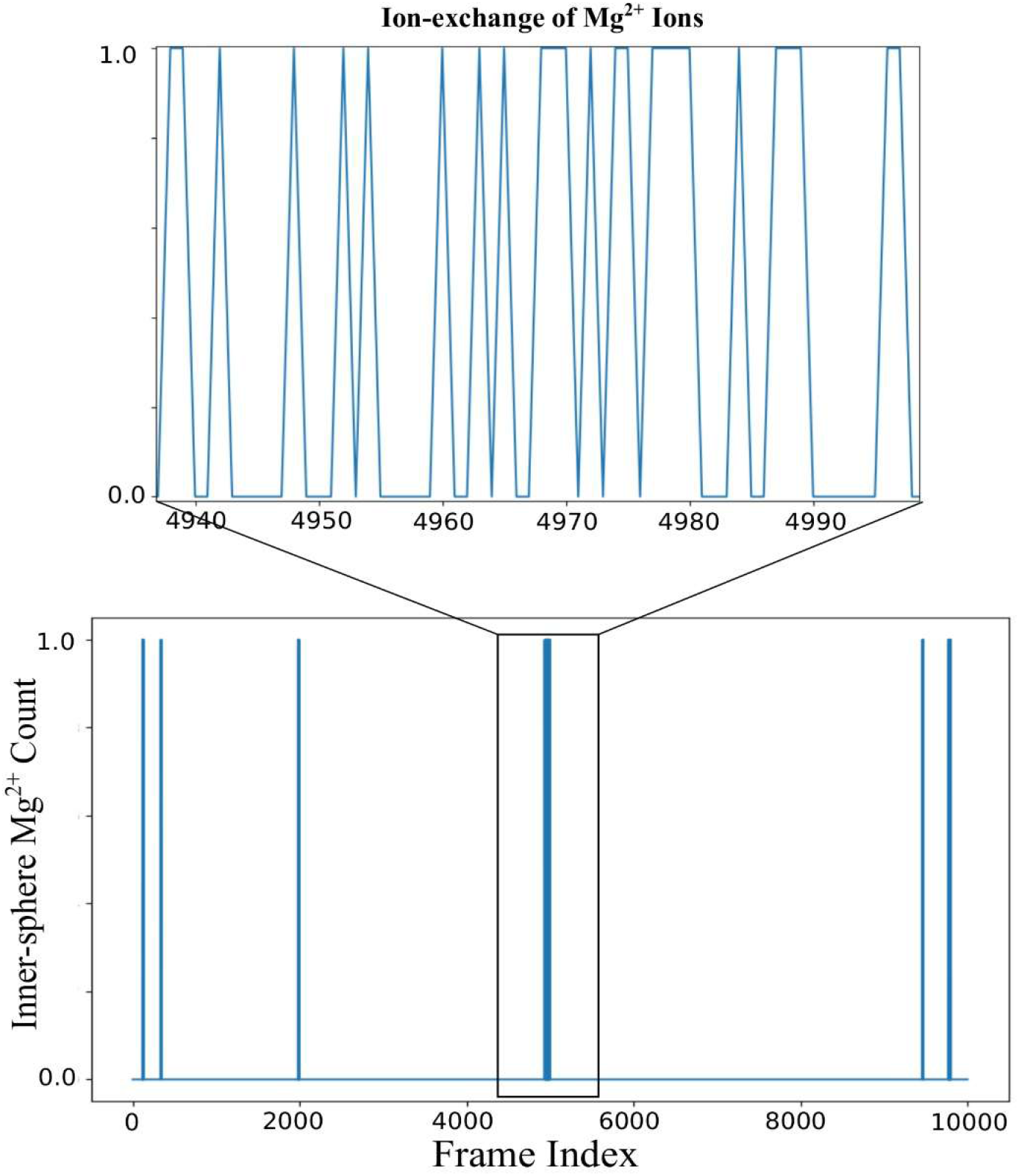
Mg^2+^ ion exchange in the flaviviral RNA system, illustrating transitions between inner- and outer-sphere coordination states.

**Figure S4:**
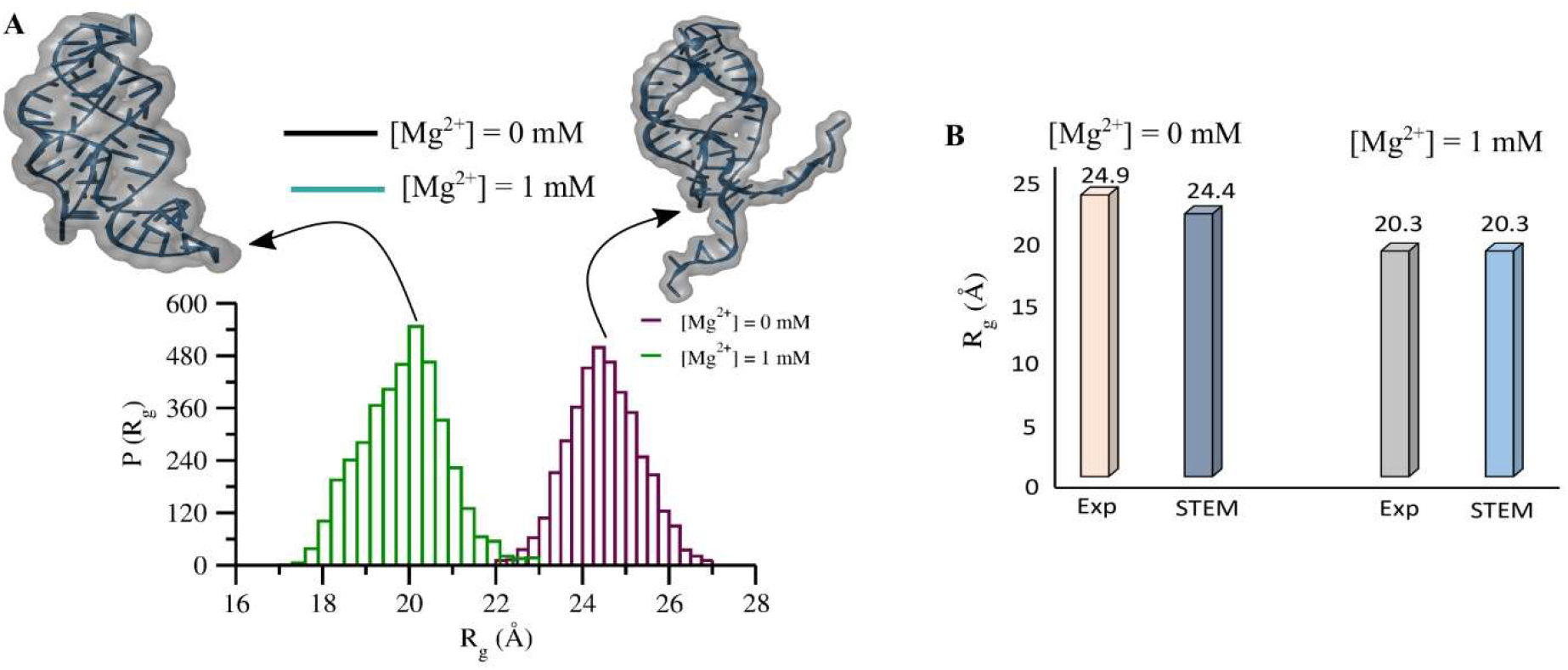
Agreement between SAXS measurements and simulations of RNA. (A) Radius of gyration (Rg) distributions for the adenine riboswitch at 50 mM KCl obtained from SAXS experiments, shown in the presence and absence of Mg^2+^, along with representative corresponding structures. (B) Representative bar plots comparing the mean Rg values from SAXS experiments and STEM simulations for the adenine riboswitch in the presence and absence of Mg^2+^.

**Figure S5:**
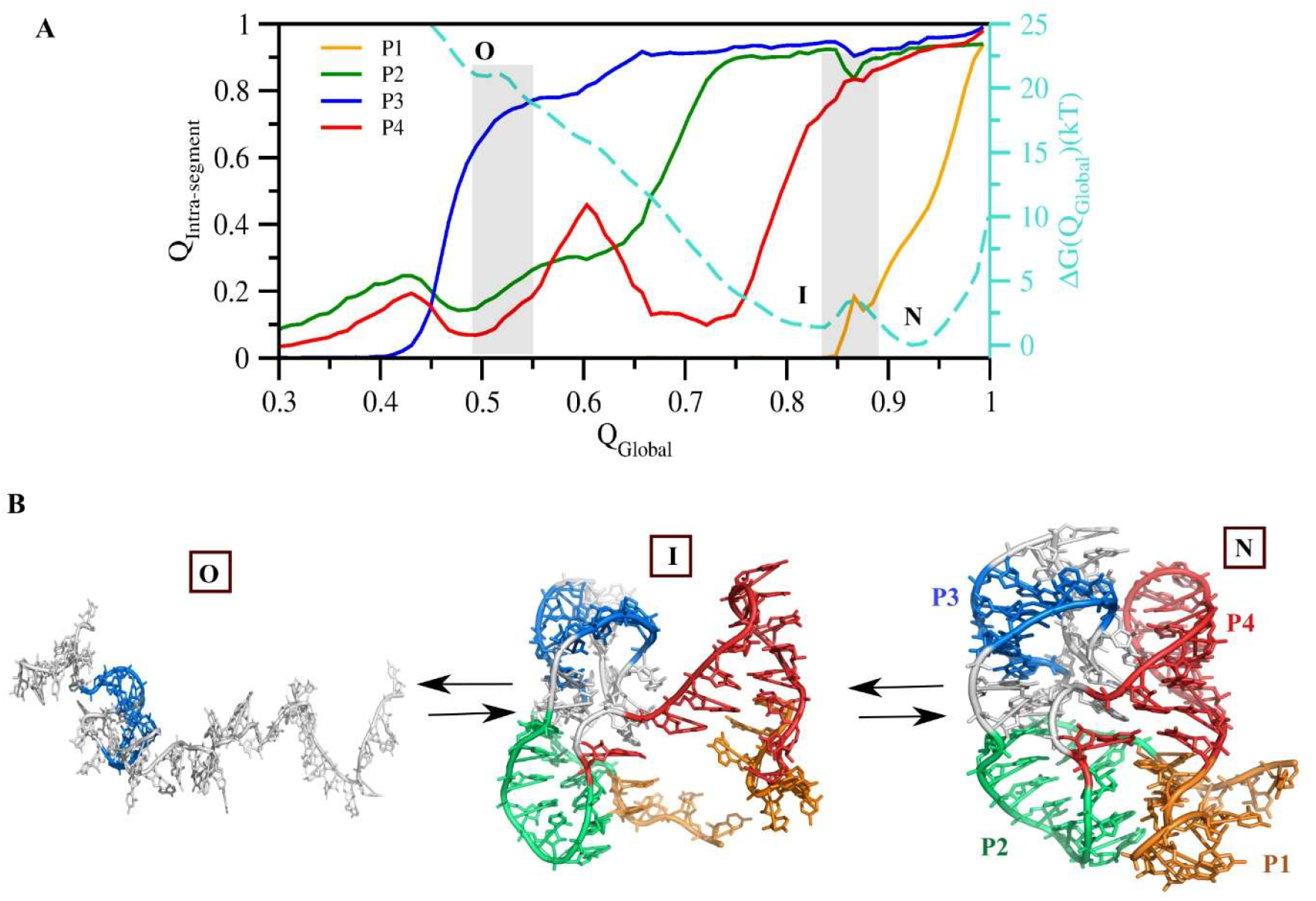
Hierarchical folding pathway of the 58-mer rRNA. (A) Sequential formation of secondary helices plotted as a function of the global fraction of native contacts (Q_Global_), where P1, P2, P3, and P4 denote the four helices. (B) Folding pathway illustrating progression from the open (O) state through the intermediate (I) state to the native (N) state.

